# Phenome-wide adaptive radiation drives lineage diversification in hyperdiverse Hymenoptera

**DOI:** 10.1101/2025.01.02.631126

**Authors:** Diego S. Porto, Lars Vilhelmsen, István Mikó, Sergei Tarasov

**Affiliations:** Finnish Museum of Natural History, University of Helsinki, Helsinki, Finland; Natural History Museum of Denmark, SCIENCE, University of Copenhagen, Copenhagen, Denmark; University of New Hampshire, Durham, NH, USA

## Abstract

How complex phenotypes evolve across the Tree of Life remains poorly understood because phenotypic evolution unfolds across multiple hierarchical and developmental scales. Hymenoptera (ants, wasps, and bees) represent one of the largest animal radiations (>150,000 species), yet the origin of their remarkable phenotypic diversity remains unclear. Here, we reconstruct the evolution of the hymenopteran skeletomuscular phenome and analyze its dynamics across anatomical hierarchies and developmental stages. We show that hymenopteran adaptive radiation was a phenome-wide process facilitated by evolutionary decoupling between adults and larvae and the semi-independent evolution of adult anatomical systems. Phenotypic and lineage diversification was driven not simply by key innovations or elevated phenomic rates, but by the partitioning of evolutionary innovation across anatomical systems, generating a broader repertoire of adaptive solutions. These findings extend the classical view of adaptive radiation from individual key innovations to phenome-wide reorganization.

## Introduction

“Endless forms most beautiful”, as Charles Darwin poetically described the diversity of organisms, motivates a fundamental question in biology: what generates large-scale phenotypic diversification across the Tree of Life? Among the most striking examples are the Hymenoptera, a hyperdiverse clade exhibiting extraordinary morphological, behavioural, and ecological variation^1,2^. Originating in the Permian (∼280 Ma)^3,4^, these insects have transformed terrestrial ecosystems and human societies through pollination, biological control, honey production, and cultural symbolism ^5,6^. Despite their prominence, the processes underlying their remarkable phenotypic diversification remain poorly understood.

A prevailing explanation attributes hymenopteran success to the emergence of four key innovations – parasitoidism, the wasp waist, the stinger apparatus, and the secondary reversals to phytophagy in several apocritan lineages^1,3^. Arising sequentially from the Triassic to the Palaeogene, these traits are thought to have catalyzed major hymenopteran radiations^2,3^. However, empirical support for this view is limited. Except for secondary phytophagy (present in ∼15% of species), none of these innovations have been robustly linked to lineage diversification^3^, and the relationship between morphological evolution and lineage diversification remains unclear^1,2^. Fossil and phylogenetic evidence further indicate that Hymenoptera experienced repeated pulses of diversity expansion and decline in various lineages, suggesting a more complex history than predicted by the simple key-innovation scenario^3,4^.

Besides the key innovations, another mechanism potentially driving hymenopteran diversification is associated with complete metamorphosis^7^. The adaptive decoupling hypothesis posits that the developmental separation of larval and adult stages in holometabolous insects enables selection to act independently on each life stage, promoting inter-stage niche differentiation and reducing ecological trade-offs^8,9^. However, the role of this mechanism in shaping phenotypic evolution remains largely untested in Hymenoptera^10^ and across Holometabola more broadly^8,9^.

Progress in resolving these questions has been hindered by a fundamental limitation: morphological evolution is typically studied using a small number of traits^11,12^, despite phenotypes being inherently multi-dimensional and hierarchically organized^13,14^. In holometabolous insects, this complexity is further amplified by complete metamorphosis, which introduces an additional developmental dimension through the decoupling of larval and adult phenotypes. Furthermore, anatomical systems vary in their degree of developmental and functional integration, forming semi-independent modules that evolve across multiple organizational scales ^11,15–17^. Compressing this multifaceted complexity into a limited set of traits may therefore obscure the mechanisms governing phenotypic diversification and adaptive radiation^13,18,19^.

Here, we reconstruct the dynamics of phenotypic diversification associated with major structural transitions in Hymenoptera across hierarchical anatomical and developmental scales. To achieve this, we employ phenomic units – higher-order ensembles of discrete traits that capture integrated evolution of anatomical systems^13^. Rather than treating individual traits as independent evolutionary units^20^, our framework considers anatomical systems as the fundamental units of evolution (Fig. 1a), enabling their dynamics to be reconstructed across multiple levels of biological organization.

**Figure 1.**
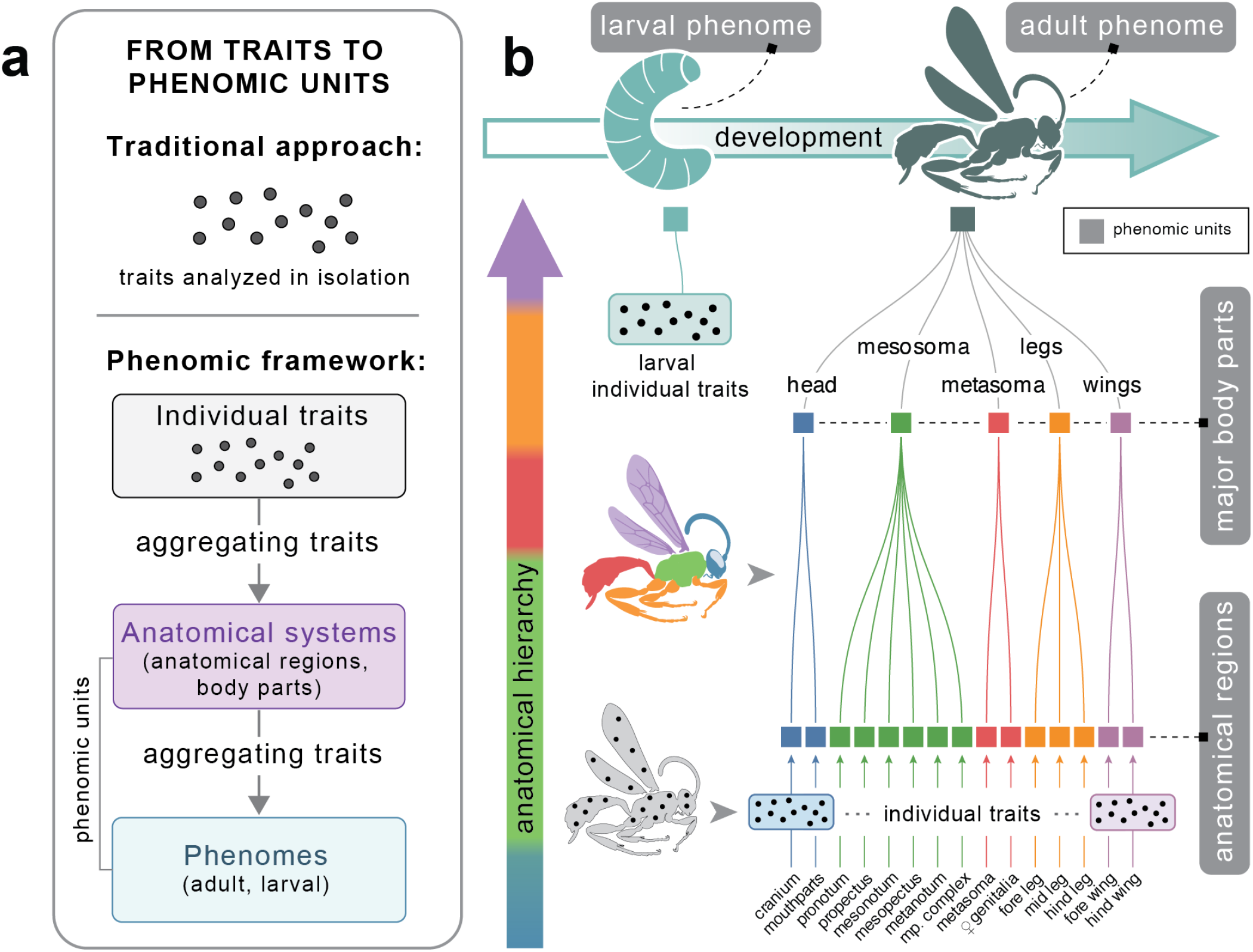
Phenomic units and hierarchical organization of the phenome. **a**, Traditional traitbased approaches treat individual traits as isolated evolutionary units. In contrast, the phenomic approach aggregates anatomically related traits into higher-order phenomic units, enabling the integrated dynamics of anatomical systems and phenomes to be reconstructed. **b**, Distribution of the phenomic units analyzed in this study across hierarchical levels of anatomical organization (15 anatomical regions, 5 major body parts, and the adult phenome) and developmental stages (larva and adult phenomes).

To investigate the evolution of phenomic units, we applied this framework to a dataset of 383 discrete traits^21,22^ describing the adult skeletomuscular system across 77 species representing major hymenopteran lineages. The skeletomuscular system was chosen because it underpins the functional and ecological strategies that enable insects to adapt and diversify ^23^. Discrete traits are especially suited to modelling skeletomuscular system because they explicitly capture the gain, loss, and transformation of anatomical structures – processes fundamental to phenotypic evolution at large timescales^24,25^.

Using ontology-informed reconstructions^13^, we analyzed the evolution of phenomic units across a phylogenomic timetree^3^ and three levels of anatomical organization (Fig. 1b): (i) anatomical regions; (ii) major body parts; and (iii) the integrated adult phenome. This approach allowed us to assess how evolutionary change is distributed across anatomical scales, distinguishing localized modifications from coordinated, phenome-wide transformations. To investigate evolution along the developmental axis and test the adaptive decoupling hypothesis, we additionally reconstructed larval phenome evolution (Fig. 1b).

Finally, we developed new metrics to quantify phenomic diversification through time. Together, these analyses allowed us to reconstruct the tempo and mode of phenome evolution and assess how lineage diversification, developmental decoupling, and phenotypic integration shaped major evolutionary transitions in Hymenoptera, giving rise to the extraordinary diversity of this taxon.

### Decoupled dynamics of adult and larval phenomes

Under the adaptive decoupling hypothesis, larval and adult phenotypes are expected to evolve under partially independent selective regimes and therefore exhibit weakly coupled evolutionary dynamics. Reconstructed rates for adult and larval phenomes reveal broadly similar temporal trajectories, with elevated rates during the Triassic followed by a long-term decline toward the present (Fig. 2a,b). However, the magnitude and timing of these accelerations differ between life stages: adult morphological evolution peaks in the Late Triassic (∼226 Ma) at 4.7× the mean rate, whereas larval evolution peaks earlier, in the Early Triassic (∼247 Ma), at 2.5× the mean rate.

**Figure 2.**
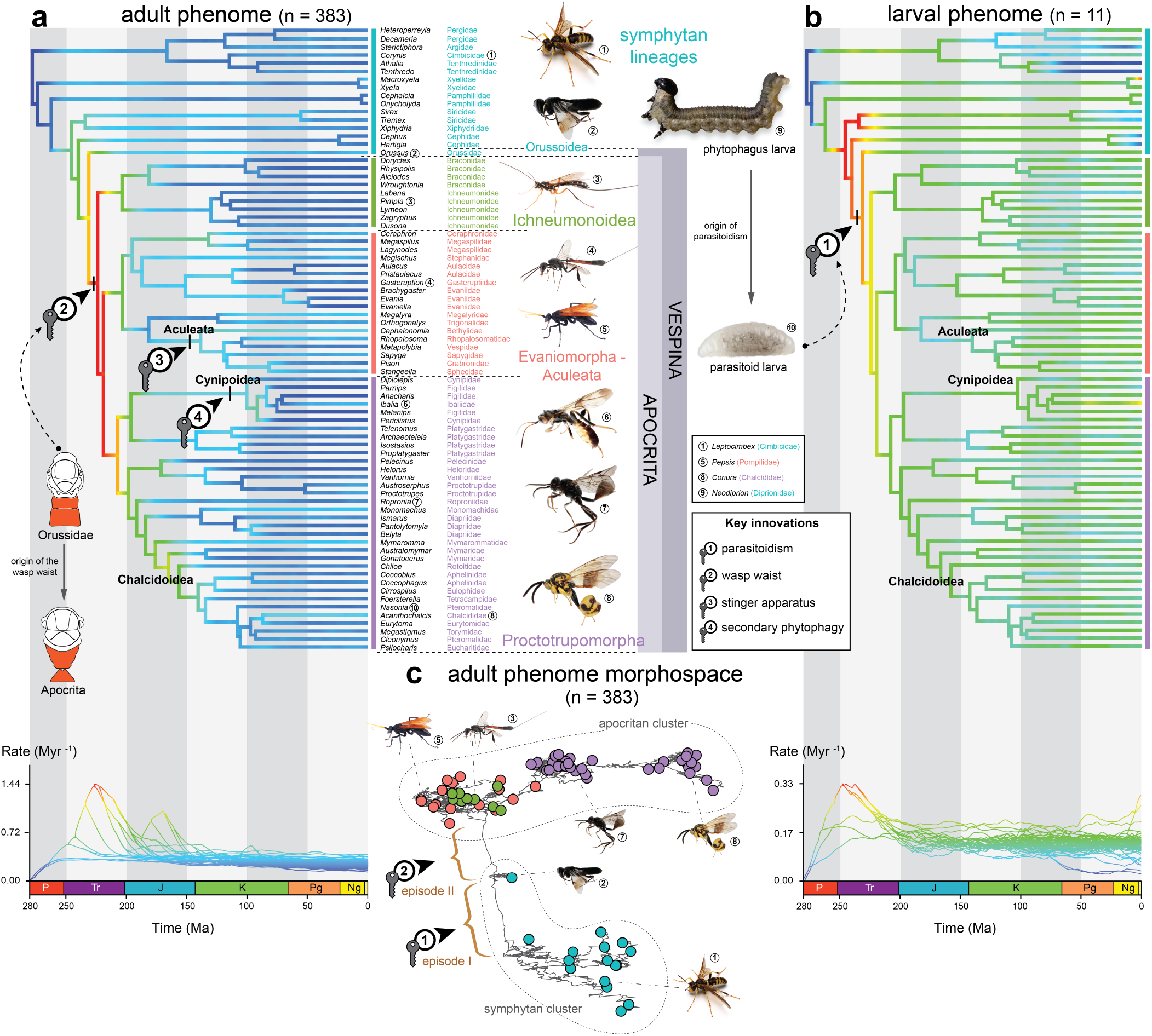
Key innovations and evolutionary dynamics of adult and larval phenomes in Hymenoptera. **a**, Evolutionary rates inferred for the adult phenome (rate profile shown below) peak near the origin of the wasp waist and decline toward the present. **b**, Evolutionary rates inferred for the larval phenome (rate profile shown below) peak shortly before the transition from phytophagy to larval parasitoidism and subsequently decline toward the present. **c**, Adult phenome morphospace reveals two major phenotypic leaps, associated with the origin of larval parasitoidism (episode I) and the origin of the wasp waist (episode II). Values in parentheses indicate number of discrete traits comprising each phenomic unit. Image credits to David Cheung (5), Hiroshi Nakamine (1), and Line Kræmer (3,4,9).

To test whether these temporal similarities reflect coupled evolutionary dynamics, we compared branch-specific rate estimates between life stages using Pearson’s correlation coefficient (*r*). Despite their similar dynamics, adult and larval rates are not correlated (*p* = 0.43). Moreover, none of the 15 adult anatomical regions show significant associations with larval rates (all *p* ≫ 0.05; Fig. 3b). These results indicate evolutionary independence between larval and adult stages, providing macroevolutionary support for the adaptive decoupling hypothesis in Hymenoptera. Although developmental decoupling has previously been documented in vertebrates^26^, comparable evidence from Holometabola has been limited. Our study provides the first phenome-wide demonstration of this phenomenon in a holometabolan order, motivating similar analyses across other insects.

**Figure 3.**
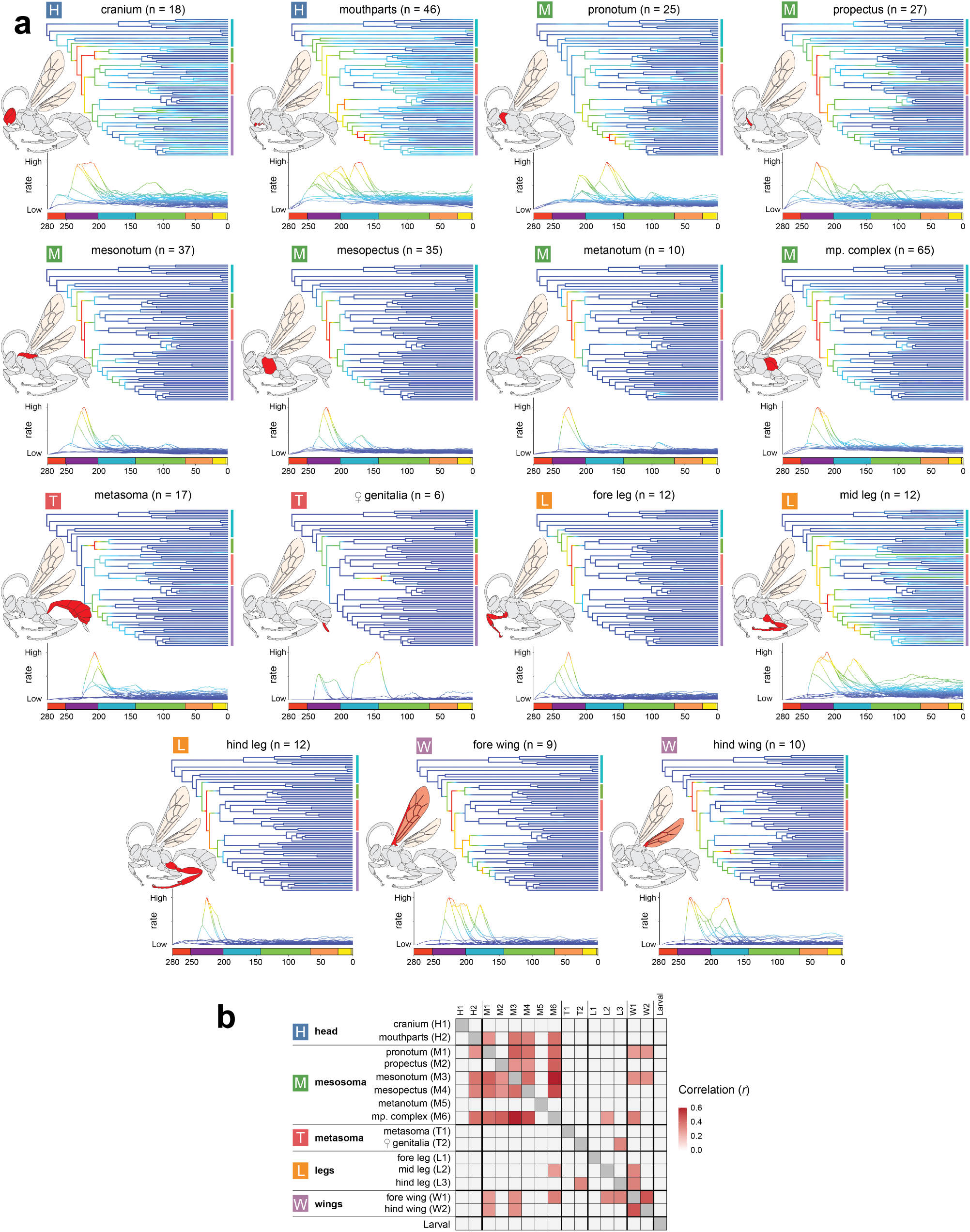
Inferred evolutionary rates across anatomical regions and their integration. **a**, Anatomical regions exhibit heterogeneous patterns of rate variation across lineages and through time. Values in parentheses indicate number of discrete traits comprising each phenomic unit. **b**, Pairwise correlations of evolutionary rates (Pearson’s *r*) reveal largely independent dynamics between the larval phenome and adult anatomical regions, whereas adult anatomical regions exhibit predominantly semi-independent dynamics. The mesosoma is an exception, forming a more tightly integrated evolutionary module with stronger correlations among its constituent regions. Only statistically significant correlations (*p <* 0.05) are shown.

Notably, the peak evolutionary rates in the larval and adult phenomes coincide with the emergence of two key innovations: larval parasitoidism in Vespina and the wasp waist in Apocrita (Fig. 2a,b). The larval innovation involved a shift from a caterpillar-like form – with elongated bodies and well-developed appendages – to a compact, grub-like morphology with reduced appendages, reflecting a transition from herbivory to parasitoidism^2,27^. The wasp waist evolved through the fusion of the first abdominal segment with the thorax in the adult, forming two distinct new body tagmata (Extended Data Fig. 1): the mesosoma (thorax plus first abdominal segment) and metasoma (remaining abdomen). This reorganization increased metasomal mobility and improved oviposition, facilitating the parasitoid lifestyle in Apocrita ^2,28^.

The reconstructed morphospace reveals that the origin of the wasp waist represents a major transition in the adult skeletomuscular system, separating Hymenoptera into two distinct clusters (Fig. 2c): a smaller, more morphologically conservative symphytan cluster (5.2% of extant species) and a large apocritan cluster (94.8% of extant species). This transition was not abrupt but unfolded gradually over ∼25 Ma during the Triassic through two successive episodes of phenome-wide reorganization, hereafter referred to collectively as the apocritan transition. The first episode coincided with the origin of the parasitoid lifestyle, during which the larval phenome also experienced its largest evolutionary reorganization, whereas the second culminated in the emergence of the wasp waist and was restricted to the adult stage (Fig. 2c). The magnitude of phenotypic change, measured using Hamming distance, was nearly identical between the two episodes (mean = 30.9, s.d. = 3.81 and mean = 30.6, s.d. = 3.42; respectively), indicating that both represented comparable phenomic transitions. These results show that major reorganizations of the larval and adult phenomes can occur either simultaneously or independently. Rather than requiring temporally separated evolution of life stages, developmental decoupling appears to permit both coordinated and stage-specific phenomic innovation, thereby facilitating rapid exploration of morphospace. We interpret these transformations as likely responses to ecological opportunities that arose following the Permian–Triassic mass extinction, when newly available niches were readily occupied by novel phenotypes ^29^.

### Adult phenome evolution is weakly integrated but episodically coordinated

Evolutionary rates reconstructed across 15 adult anatomical regions reveal an overall early-burst–like decline but differ markedly in temporal dynamics (Fig. 3a). Some anatomical regions closely track the trajectory of the adult phenome (e.g. mesonotum and mesopectus; Fig. 3a), whereas others show multiple accelerations throughout the Triassic–Jurassic interval (e.g. mouthparts and pronotum; Fig. 3a). Similar results were observed for the major body parts (Supplementary Note 1; Supplementary Fig. 2). Despite these temporal similarities, overall phenotypic integration among regions is weak (Fig. 3b). Mean absolute pairwise correlation of branch-specific rates was low (*r* = 0.084 *±* 0.16 s.d.): only 23 of 105 pairwise comparisons were statistically significant after Bonferroni correction (*p <* 0.05). Significant correlations were concentrated primarily within the mesosoma (consisting of six anatomical regions), where within-module correlations exceeded those between mesosomal and non-mesosomal regions (*r*_within_ = 0.29 versus *r*_between_ = 0.17; *p* = 0.015). Together, these results indicate that most adult anatomical regions evolve semi-independently; only the mesosoma forms a distinct evolutionary module, likely reflecting morpho-functional changes associated with the the wasp waist innovation (Fig. 3b).

To quantify temporal coordination of evolutionary change across anatomical regions, we identified regionspecific accelerated branches in the Hymenoptera phylogeny, defined as branches with exceptionally rapid rates (top 5% of the rate distribution). Such episodes often mark major macroevolutionary trends or the emergence of morphological innovations, and therefore capture biologically meaningful deviations from background dynamics^30,31^. The proportion of anatomical regions exhibiting accelerated evolution (*P*_*c*_) peaks during the apocritan transition (Extended Data Fig. 2), indicating that major phenotypic transformations in Hymenoptera emerged through transient episodes of rapid synchronous changes superimposed on an otherwise modular, semi-independent anatomical architecture.

### The phenomic basis of lineage diversification

To quantify phenomic diversification, we developed 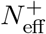 (effective number of accelerated phenomic units), which integrates the richness and evenness of accelerated anatomical regions across lineages (Fig. 4b). 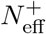 peaks near the Triassic–Jurassic boundary (∼200 Ma; Fig. 4b), shortly after the apocritan transition characterized by elevated phenomic coordination (*P*_*c*_). To test whether phenotypic diversification was associated with lineage diversification, we compared 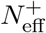 with published estimates of lineage diversification rates inferred from: (i) fossil occurrence data with PyRate^4^; and (ii) a molecular timetree with BAMM^3^. 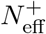 exhibited an exceptionally strong positive association with the PyRate rate in both the original and detrended analyses, indicating a robust relationship independent of long-term temporal trends (Fig. 4b; Extended Data Fig. 3a; standard: *r* = 0.95, *p* = 6.7 *×* 10^−7^, 95% CI = 0.84–0.99; detrended: *r* = 0.95, *p* = 7.9 *×* 10^−7^, 95% CI = 0.83–0.98). Moreover, both 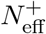 and PyRate rate show nearly identical trends when calculated over the same temporal resolution (Extended Data Fig. 3b).

**Figure 4.**
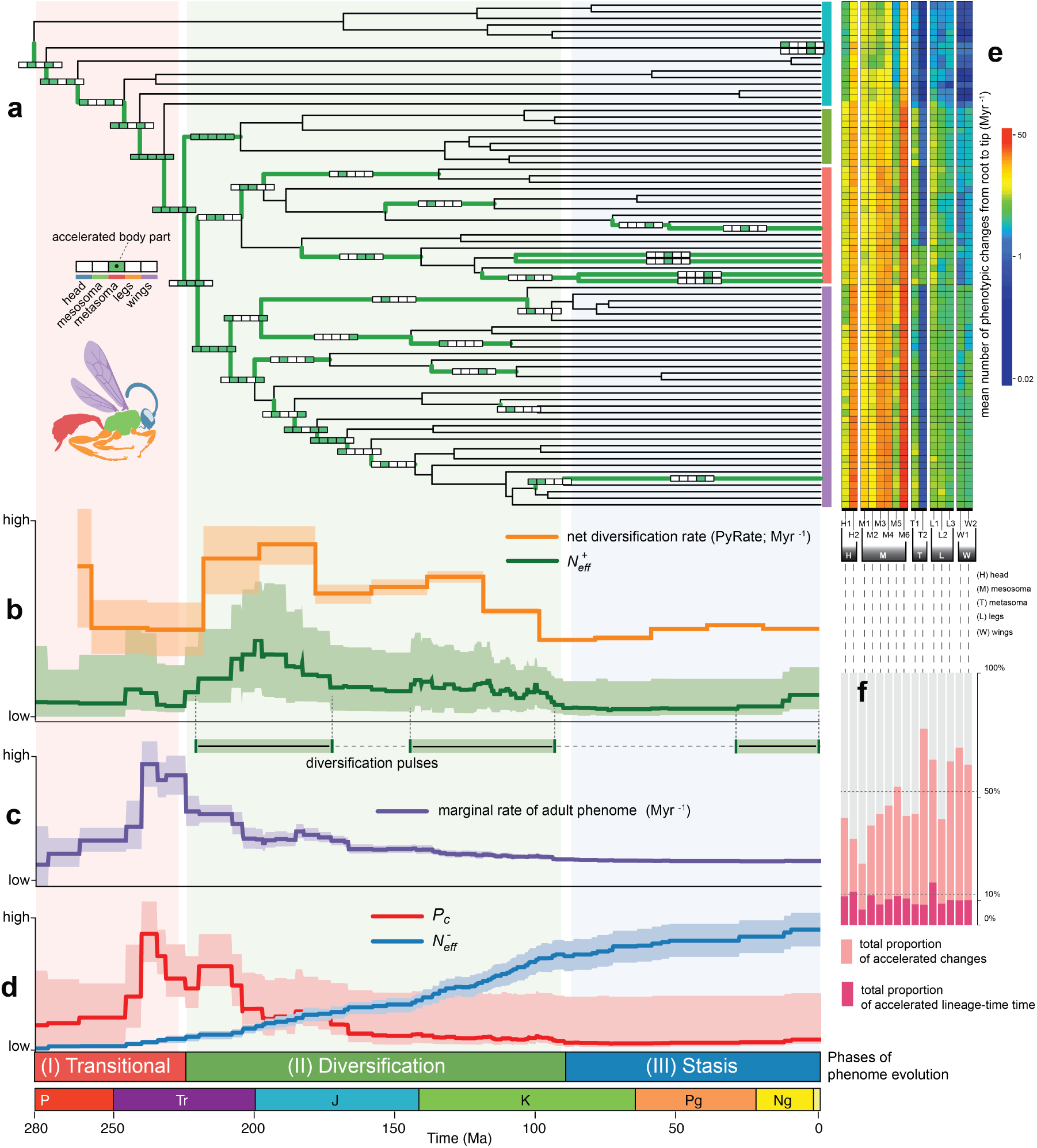
Three phases of phenome-wide adaptive radiation in Hymenoptera. **a**, Timetree showing the distribution of accelerated evolutionary episodes across lineages and major body parts. **b**, Covariation between PyRate diversification rate^4^ and the diversity of accelerated anatomical regions 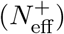, revealing coincident pulses of lineage and phenomic diversification. **c**, Peaks in the marginal rate of the adult phenome predate pulses of lineage and phenomic diversification, but are associated with temporal phenomic coordination (*P*_*c*_) shown in **(d). d**, Temporal trajectories of *P*_*c*_, the diversity of decelerated regions 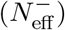 and 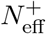 from **(b)** define three phases of hymenopteran phenome evolution. **e**, Mean number of phenotypic changes per lineage for individual anatomical regions, revealing mosaic evolution across the phenome. **f**, Proportion of total phenotypic changes and lineage time associated with accelerated episodes of adult phenome evolution. Shaded areas in panels **(b–d)** represent 95% confidence intervals, whereas the shaded area around the PyRate estimate in panel **(a)** represent the 50% credible interval.

Sensitivity analyses showed that the temporal patterns of 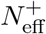 were consistent across different branch-rate percentile thresholds, demonstrating that its association with PyRate rate is robust to the choice of threshold (i.e., top 7.5%, 5% or 2.5% of the rate distribution; Supplementary Note 2; Supplementary Figs. 3-5). Moreover, decomposition of 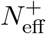 showed that this association is best explained by the combined effects of the frequency (*f* ^+^) and phenomic breadth (*H*^+^) of accelerated evolution, rather than by either component alone (Supplementary Note 3).

In contrast, BAMM rates exhibited a negative correlation with 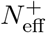 (*r* = −0.59, *p* = 0.032). However, this association disappeared after detrending (*p* = 0.6), indicating that it was driven by a shared longterm trend, namely the monotonic increase in BAMM rates through time (Supplementary Fig. 8). This discrepancy likely reflects limitations of BAMM in reconstructing deep-time diversification dynamics in the absence of fossils ^32,33^.

The strong association between 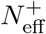 and PyRate suggests that lineage diversification in Hymenoptera was tightly coupled to the diversity of rapidly evolving anatomical regions. Phenotypic diversification was therefore a phenome-wide process associated not with rapid evolution in one or a few anatomical systems, but with the proliferation of distinct evolutionary trajectories distributed throughout the phenome (Extended Data Fig. 4). We therefore interpret elevated 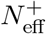 as a phenomic signature of adaptive diversification, reflecting periods when multiple anatomical systems evolved along divergent trajectories across proliferating lineages. Apparently, such episodes generated a broad spectrum of phenotypic innovations across different anatomical regions, expanding the range of phenotypic solutions available to natural selection. These innovations likely facilitated the exploitation of new ecological opportunities and increased ecological differentiation among lineages, thereby promoting lineage diversification (Extended Data Fig. 4).

The pattern of covariation between 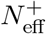 and PyRate identifies one major pulse of adaptive radiation during the Triassic–Jurassic and two smaller pulses in the Jurassic–Cretaceous and Cenozoic (Fig. 4b). At least the the first two pulses broadly coincide with periods of ecological restructuring and expansion of major hymenopteran life-history strategies^3,4^.

Marginal rate of adult phenome (Fig. 4c) shows no association with PyRate estimates (*p* = 0.56), indicating that overall morphological rate is decoupled from lineage diversification. Instead, marginal rate is positively associated with temporal phenomic coordination (*P*_*c*_; Fig. 4d; standard: *r* = 0.97, 95% CI: 0.89–0.99, *p* = 6.9 *×* 10^−8^; detrended: *r* = 0.90, 95% CI: 0.69–0.97, *p* = 2.6 *×* 10^−5^), suggesting that elevated whole-phenome rate predicts rapid, synchronous changes distributed across multiple anatomical regions. Consequently, major evolutionary transitions associated with elevated rates (e.g., apocritan transition), represent episodes of rapid phenome-wide restructuring without immediate lineage diversification.

This pattern is initially surprising because evolutionary theory predicts that rapid morphological evolution should promote lineage divergence and speciation^34,35^. One possible explanation is that such rapid restructuring may represent episodes of strong directional selection associated with shifts between adaptive peaks. During these episodes, lineages may pass through an adaptive valley where intermediate phenotypes experience reduced fitness, resulting in elevated extinction despite extensive phenotypic divergence ^36^. Our results are consistent with this interpretation. The origins of larval parasitoidism and the wasp waist coincide with elevated extinction rates inferred by PyRate^4^, offsetting lineage origination and resulting in near-zero net diversification (Extended Data Fig. 5). Thus, we interpret them as markers of an adaptive peak shift, with the intervening transition exhibiting the elevated extinction expected during traversal of an adaptive valley. This quantitatively supports the hypothesis that both innovations initially functioned as preadaptations facilitating entry into a new adaptive zone^2,27^, with major phenotypic and lineage diversification occurring only subsequently.

### Three phases of hymenopteran phenome evolution

Temporal trajectories of phenomic coordination (*P*_*c*_), accelerated diversification 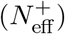, and evolutionary stasis 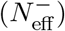 reveal three major phases of hymenopteran phenotypic evolution (Fig. 4d). Analogously to 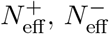 quantifies the across-lineage diversity of anatomical regions experiencing decelerated evolution.

The first phase (Permian–Late Triassic, 280–225 Ma) starts with the origin of crown Hymenoptera and culminates in the apocritan transition – an expansion from an ancestral symphytan adaptive peak to a novel apocritan peak. It involved extensive phenome-wide restructuring (elevated *P*_*c*_) associated with the evolutionary decoupling between adult and larval phenomes.

The second phase (Late Triassic–Late Cretaceous, 225–90 Ma), characterized by elevated 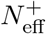, marks a period of widespread adaptive diversification across apocritan lineages. What began with the evolution of the parasitoid lifestyle ultimately gave rise to repeated ecological expansion into diverse niches, including secondary phytophagy^2,3^. This radiation was driven by the proliferation of diverse phenotypic trajectories across anatomical systems, facilitated by the semi-independent evolution of the adult phenome.

The third phase (Late Cretaceous–present, 90–0 Ma), characterized by elevated 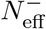, marks a period of progressive stabilization of hymenopteran skeletomuscular architecture accompanied by a slowdown in lineage diversification.

To assess how these phases contributed to present-day diversity, we quantified the mean number of phenotypic changes accumulated along each lineage from the root to the tips of the phylogeny (Fig.4e). Phenotypic change accumulated unevenly across anatomical regions and lineages, producing the mosaic pattern of divergence that characterizes extant Hymenoptera. Remarkably, ∼50% of all phenotypic changes (24–78% depending on the anatomical region; Fig.4f) occurred during episodes of accelerated evolutionary change (i.e., Phases I and II), despite these intervals comprising only ∼10% of total lineage time (6–17% depending on the anatomical region; Fig.4f). Together, these results suggest that extant hymenopteran diversity emerged through the interplay of brief bursts of phenome-wide innovation, primarily associated with the rise of Apocrita (Extended Data Fig. 6), and prolonged periods of phenotypic refinement.

## Conclusion

Phenotypic and lineage diversification in Hymenoptera is governed not simply by overall phenomic rates or isolated key innovations, but by lineages exploring distinct phenotypic trajectories through the partitioning of evolutionary innovation among anatomical systems 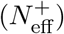. This process expands the repertoire of adaptive solutions beyond what is possible when evolution repeatedly targets the same anatomical structures. In Hymenoptera, both life stages and most anatomical regions of the adult evolve semi-independently, whereas major evolutionary transitions involve transient episodes of coordinated phenome-wide restructuring. Taken together, these results indicate that large-scale adaptive radiations may emerge through the interplay of developmental decoupling, phenotypic semi-independence, and the hierarchical organization of complex phenotypes.

Our findings are aligned with the classical Simpsonian model of adaptive radiation, in which entry into a new adaptive zone is followed by rapid phenotypic evolution, diversification, and eventual evolutionary slowdown^19^. While this pattern is widely recognized^37,38^, it has generally been inferred from variation in a small number of ecologically important traits, such as beak morphology (Darwin’s finches), ecomorphs (*Anolis* lizards), and jaw morphology (East African cichlids). Our results extend Simpson’s model by showing that adaptive radiation can operate across entire phenomes, spanning different life stages. In Hymenoptera, the transition into a new adaptive zone, its subsequent diversification, and eventual stasis (Phases I–III; Fig. 4) involved evolutionary change distributed across virtually all major anatomical systems. Thus, over macroevolutionary timescales, adaptive radiation may be driven by phenome-wide reorganization rather than the diversification of a few ecologically important traits.

This study may help explain why associations between key innovations and lineage diversification, as well as between morphological evolutionary rates and lineage diversification, are often weak or absent across diverse organisms ^3,30,31,39–41^ Neither key innovations nor elevated morphological rates alone appear sufficient to explain the phenome-wide evolutionary dynamics. Instead, these dynamics may be better captured by how phenotypic acceleration is partitioned across anatomical systems 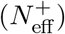. Therefore, focusing on a handful of conspicuous key innovations provides an incomplete view of macroevolution. Phenomic units overcome this limitation by revealing dynamics that remain largely hidden from single-trait approaches.

However, several limitations still remain to our approach. Our analyses focused on major skeletomuscular transformations, not capturing the full complexity of hymenopteran phenotypes (e.g., physiology, behaviour). Moreover, larval phenomes were reconstructed as single integrated units rather than hierarchical systems. Future incorporation of additional trait classes, hierarchical larval datasets, and fossils may provide a more nuanced view of hymenopteran evolution. Integrating phenomic and genomic data^42^ may ultimately reveal how genotype–phenotype interactions drive macroevolutionary dynamics across deep time.

This study raises a broader question: how common are phenome-wide adaptive radiations across the Tree of Life? Our framework provides a general approach for reconstructing phenome evolution and identifying the phenomic signatures of adaptive diversification. It also highlights the untapped potential of the large morphological matrices accumulated over decades for phylogenetic research^43^ as a pivotal resource for phenomic studies.

## Methods

### Morphological Data

The adult dataset used in the phenomic analyses comprised 383 characters filtered, verified, and modified from^21^ (Supplementary Methods; Supplementary Fig. 1). The larval dataset comprised 11 characters modified from^22^. Because *>*90% of Hymenoptera have larvae with highly reduced or poorly documented morphology owing to their parasitic and cryptic lifestyles respectively^6^, only a limited number of larval traits can be scored consistently across the phylogeny^2,22^ (Supplementary Methods).

To assess whether the morphological dataset captures genuine patterns of evolutionary change rather than an artefact of character selection for higher-level systematics, we evaluated its correspondence with molecular branch lengths and clade age (Supplementary Note 4; Supplementary Figs. 6,7).

### Reconstructing evolution of phenomic units

Evolution of phenomic units was reconstructed using ontophylo ^13^. The method reconstructs ancestral states of multiple discrete traits via stochastic mapping and subsequently amalgamates individual stochastic maps into composite histories representing the evolution of phenomic units. These amalgamations are enabled by annotating traits with the Hymenoptera Anatomy Ontology ^44^, allowing developmental dependencies among traits to be accounted for (e.g., wing venation cannot evolve if the wing is absent) and enabling reconstructions across multiple levels of the anatomical hierarchy ^14,25,45^ (Fig. 1). Evolutionary models were built and fitted using rphenoscate^45^ and corHMM^46^ and stochastic maps were reconstructed on a dated phylogeny modified from topology C1 of^3^ (pruned to 77 taxa) using phytools^20^. For each trait, we fitted a set of candidate models and selected the best-fitting model using the Akaike Information Criterion (Supplementary Methods).

The resulting stochastic maps were translated into three classes of evolutionary-rate estimates for downstream analyses: (i) continuously varying rates, allowing rate variation within branches and inferred under a non-homogeneous Poisson process^45^; (ii) branch-specific rates, assumed constant along individual branches; and (iii) marginal rates, representing time-dependent averages across lineages. Continuously varying rates were inferred from a subsample of 100 stochastic-mapping replicates and were used to visualize rate dynamics within branches and across the phylogeny. Branch-specific (constant) and marginal rates were estimated from 1,000 stochastic-mapping replicates. For each replicate, rate-through-time trajectories were calculated by interpolating evolutionary rates across 1,000 evenly spaced time points (Supplementary Methods). Branch-specific (constant) and marginal rates were subsequently used to assess rate correlations among anatomical regions and associations with phenomic metrics.

### Morphospace reconstruction

The morphospace was reconstructed in ontophylo from a stochastic map of the adult phenome (n = 383 discrete traits). Hamming distances among phenomic states were projected into a low-dimensional space using multidimensional scaling (Supplementary Methods).

### Correlations between branch-specific evolutionary rates

We used reconstructed mean branch-specific evolutionary rates to examine rate correlations between (i) adult and larval morphologies, and (ii) each of 15 adult elementary anatomical regions and the larval characters (Fig. 1). Following the approach used for testing gene correlations^47^, we quantified covariation using Pearson correlation coefficient (*r*). To improve normality, rates were log-transformed and then Z-transformed. Outliers were removed using a 1–99% quantile filter, and *p*-values were adjusted for multiple comparisons using a Bonferroni correction.

### Testing mesosoma modularity

To evaluate whether the six anatomical regions constituting the core-mesosoma (i.e., excluding legs and wings) form an independent evolutionary module, we performed a permutation test using the pairwise evolutionary-rate correlation matrix obtained above. From the observed matrix, we calculated (i) all correlations among the six core-mesosomal regions (i.e. within-module: pronotum, propectus, mesonotum, mesopectus, metanotum, and metapectal-propodeal complex), and (ii) all correlations between core-mesosomal regions and other anatomical regions (i.e. between-module). We then computed the observed difference 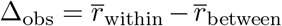, where *r* denotes the absolute mean Pearson correlation. To test whether this difference exceeded expectations under the null hypothesis of no modular structure, we permuted trait labels while preserving the number of traits assigned to the core-mesosoma module. For each permutation, we recalculated the within- and between-module correlations and obtained a null distribution of Δ values. The empirical two-tailed *p*-value was calculated as the proportion of permuted values for which |Δ| ≥ |Δ_obs_|.

### Identifying accelerated and decelerated evolutionary regimes

For each anatomical region, we quantified the empirical distribution of branch-specific rates across 1,000 stochastic-map replicates and identified unusually high and low rates using the 95th and 5th percentiles of the branch-rate distribution as thresholds, respectively. Branches with rates above the 95th percentile were classified as accelerated, whereas those below the 5th percentile were classified as decelerated.

To summarize patterns across the Hymenoptera phylogeny, we used a binary coding scheme in which enriched branches (accelerated or decelerated) were assigned a value of 1 and all other branches a value of 0. Major body parts (head, mesosoma, metasoma, legs, and wings) were considered enriched if any of their constituent anatomical regions were enriched.

### Estimating phenomic coordination (*P*_*c*_)

Phenomic coordination (*P*_*c*_) quantifies the extent to which accelerated evolutionary regimes are coordinated across branches and organismal anatomy. For each branch of the phylogeny, we calculated the proportion of anatomical regions classified as accelerated (i.e. assigned a value of 1 under the binary coding scheme described above). This proportion measures how broadly accelerated evolution is distributed across the phenome along a given branch. We then estimated *P*_*c*_ by averaging these proportions across all branches within each of 1,000 evenly spaced time slices spanning the phylogeny. Uncertainty was quantified using empirical 95% confidence interval. The resulting metric, *P*_*c*_, reflects the average fraction of anatomical regions evolving in an accelerated regime along time. Higher values indicate greater phenomic coordination, with accelerated evolution occurring simultaneously across a larger proportion of the organismal anatomy.

### Estimating phenomic diversity metrics (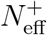 and 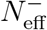)

To quantify the temporal diversity of phenomic units evolving in accelerated or decelerated regimes, we developed the metrics 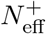 (effective number of accelerated phenomic units) and 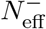 (effective number of decelerated phenomic units). These metrics are conceptually analogous to the effective number of species used in community ecology derived from Shannon entropy^48^. They summarize the diversity of accelerated or decelerated phenomic units by accounting for both their richness and their evenness of occurrence across lineages, thereby providing a single measure of phenomic unit diversity through time (Extended Data Fig. 4).

The metrics were calculated independently for accelerated and decelerated evolutionary regimes. At each time slice *t* of a tree, every branch was represented by a binary vector of length *L*, corresponding to the elementary phenomic units (*L* = 15), where a value of 1 indicated an accelerated (or decelerated) phenomic unit and 0 otherwise. A time slice consisted of *N* such branch vectors, where *N* is the number of branches intersecting that slice. Let *N* ^*±*^ denote the number of branches containing at least one accelerated (+) or decelerated (−) phenomic unit, and let 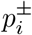 denote the frequency of each distinct binary pattern among these branches. The Shannon entropy of the distribution of patterns is

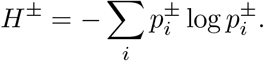

The effective number of phenomic units is then defined as

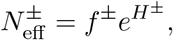

where

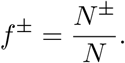

The quantity *f* ^*±*^ measures the frequency of accelerated (+) or decelerated (−) evolution by quantifying the proportion of branches exhibiting at least one accelerated or decelerated phenomic unit. The quantity *H*^*±*^ is the corresponding Shannon entropy and measures the phenomic breadth of accelerated or decelerated evolution across anatomical regions. By construction, 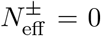 when no accelerated (or decelerated) phenomic units are present, and larger values indicate greater diversity of phenomic-unit patterns across lineages (Extended Data Fig. 4).

Temporal trajectories of 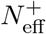 and 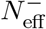 were estimated across 1,000 stochastic-map replicated. For each replicate, the metrics were calculated independently over 1,000 evenly spaced time slices spanning the phylogeny. Mean trajectories were then obtained by averaging across stochastic-map replicates. Uncertainty was quantified using empirical 95% confidence interval.

### Correlations between phenomic metrics and evolutionary rates

PyRate diversification rates were obtained from the posterior sample of the Birth–Death model with constrained shifts and without singletons (BDCS-fam-ssing-20)^4^. We selected this analysis because it most closely matches our study, with both estimating diversification at the family level across Hymenoptera; all analyses were restricted to the interval spanning the present to the oldest confirmed Hymenoptera fossil occurrence^4^. To ensure direct comparability, all metrics and rates were averaged within the 20-million-year time bins used to estimate PyRate diversification rates in the original study.

We examined four relationships: (i) PyRate diversification rate versus 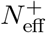; (ii) BAMM diversification rate^3^ versus 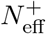; (iii) PyRate diversification rate versus adult phenomic evolutionary rate; and (iv) phenomic coordination (*P*_*c*_) versus adult phenomic evolutionary rate.

Because temporal series may share long-term trends that can inflate correlation estimates, we performed both standard and detrended Pearson correlation analyses. Detrended correlations were calculated from residuals obtained by regressing each variable against time, thereby testing associations independent of shared temporal trends.

### Estimating phenotypic change accumulated at the tips

For each of 15 adult anatomical regions, stochastic maps reconstructed with ontophylo were used to calculate the mean number of phenotypic changes accumulated from the root to each tip across 1,000 stochasticmapping replicates. To quantify the contribution of accelerated evolutionary episodes, we calculated the proportion of root-to-tip phenotypic changes occurring along accelerated branches relative to the total number of changes accumulated along each lineage.

## Data availability

All data produced, modified, and analysed in the present study are available on GitHub (https://github.com/diegosasso/Hymenoptera_paper) and Zenodo (link to be added).

## Code availability

The code necessary to replicate the analyses and figures of the present study are available on GitHub (https://github.com/diegosasso/Hymenoptera_paper) and Zenodo (link to be added).

## Acknowledgements

D.S.P. was supported by the Research Council of Finland (grant 362624). S.T. was also supported by the Research Council of Finland (grants 361983, 346294, 339576 and 369299). We thank Dr. Corentin Jouault for kindly sharing the PyRate data of Hymenoptera. We are grateful to Dimitar Dimitrov, Josef Uyeda and Matthew Pennell for valuable comments that helped improving earlier versions of this manuscript.

## Author contributions

D.S.P. and S.T. conceptualized and designed the study, analysed the data, and wrote the first draft of the manuscript. D.S.P., I.M., and L.V. curated the data. All authors contributed to the final draft and revisions of the manuscript.

## Additional information

### Supplementary information

The online version contains Supplementary Information available at link to be adde

**Correspondence and requests for materials** should be addressed to D.S.Porto and S.Tarasov.

## Extended Data

**Extended Data Figure 1.**
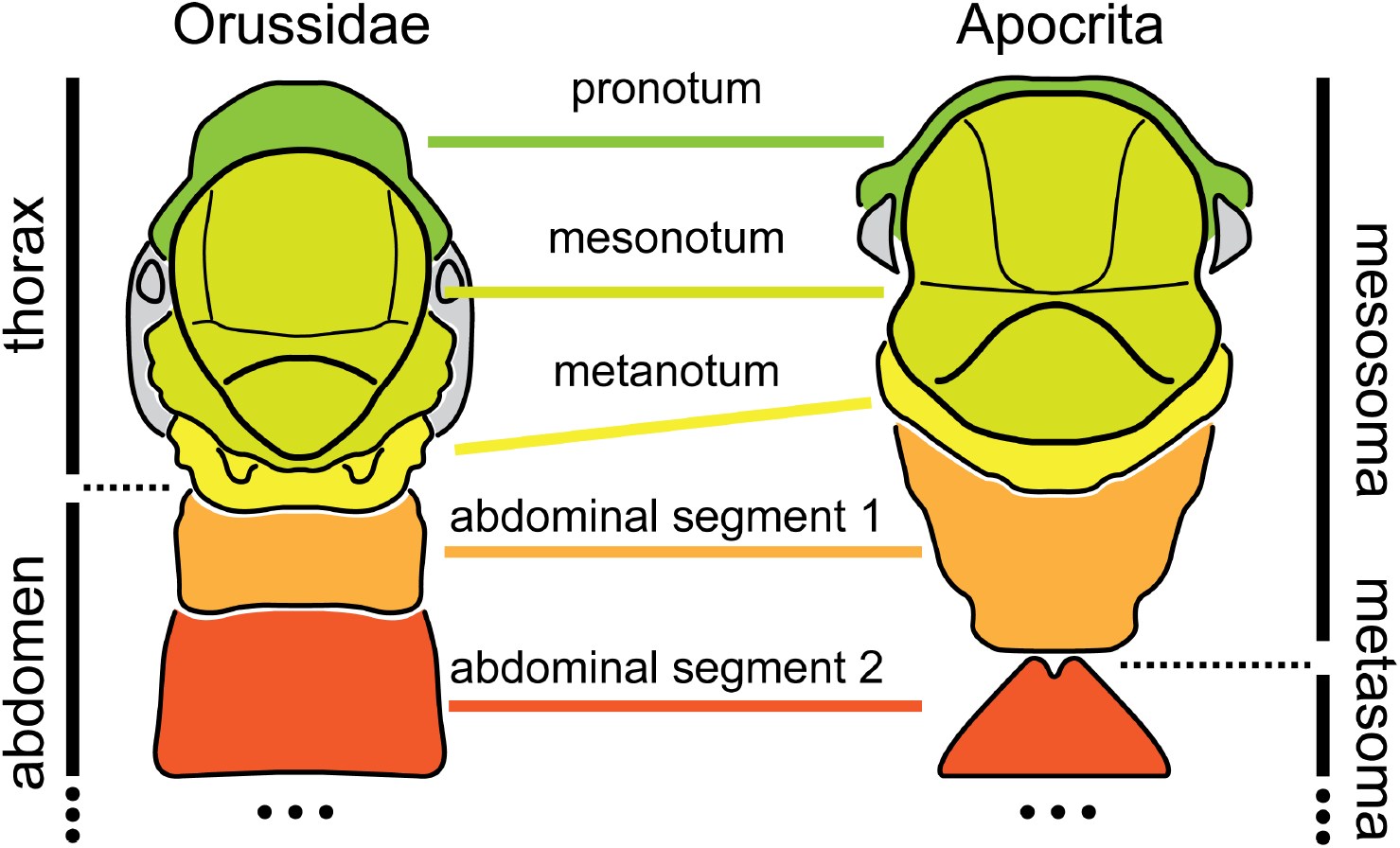
Wasp waist innovation. Anatomical diagrams of adult Hymenoptera illustrating the conditions of the thorax and abdomen in dorsal view before (left) and after (right) the wasp waist transformation.

**Extended Data Figure 2.**
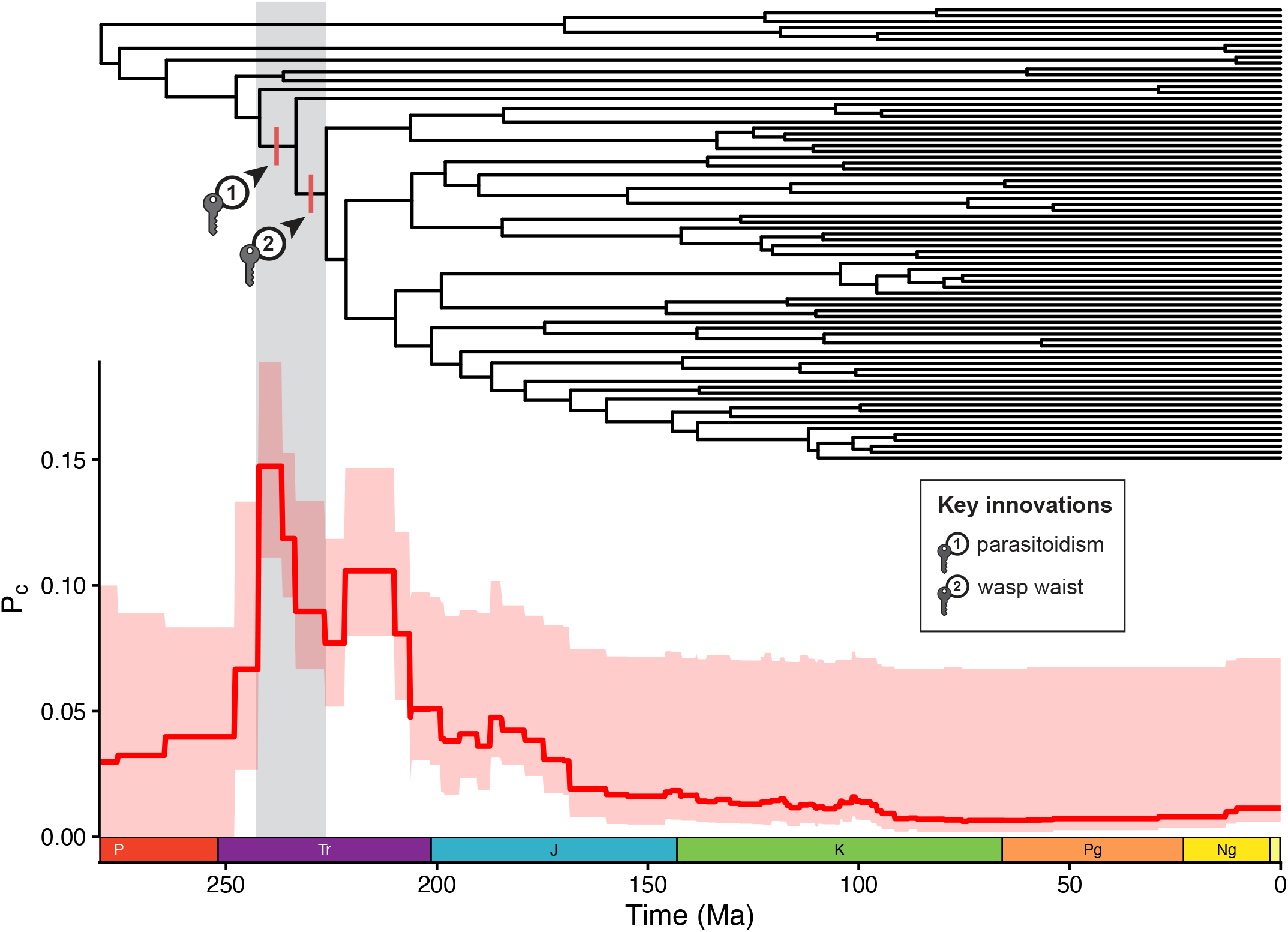
Key innovations coincide with elevated phenomic coordination (*P*_*c*_). This pattern indicates that the origins of larval parasitoidism and the adult wasp waist, which together mark a major restructuring of the hymenopteran body plan, coincided with a period when accelerated evolutionary rates were distributed across most anatomical regions of the phenome.

**Extended Data Figure 3.**
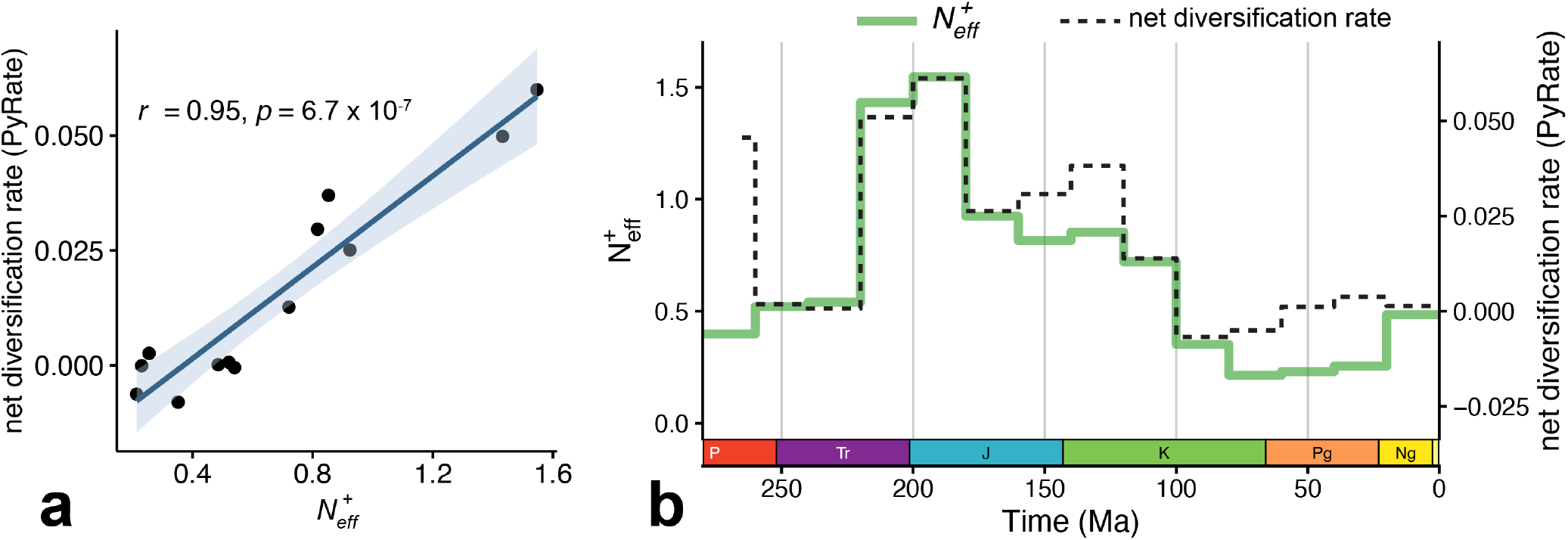
Relationship between 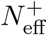 and net diversification rate inferred with PyRate^**4**^. **a**, Linear regression showing a significant positive association between 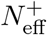 and net diversification rate. **b**, Closely matching temporal trajectories of 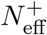 and net diversification. 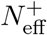 estimates from Fig. 4b were averaged within the same 20-Myr time bins used in the original PyRate analysis to ensure a common temporal resolution.

**Extended Data Figure 4.**
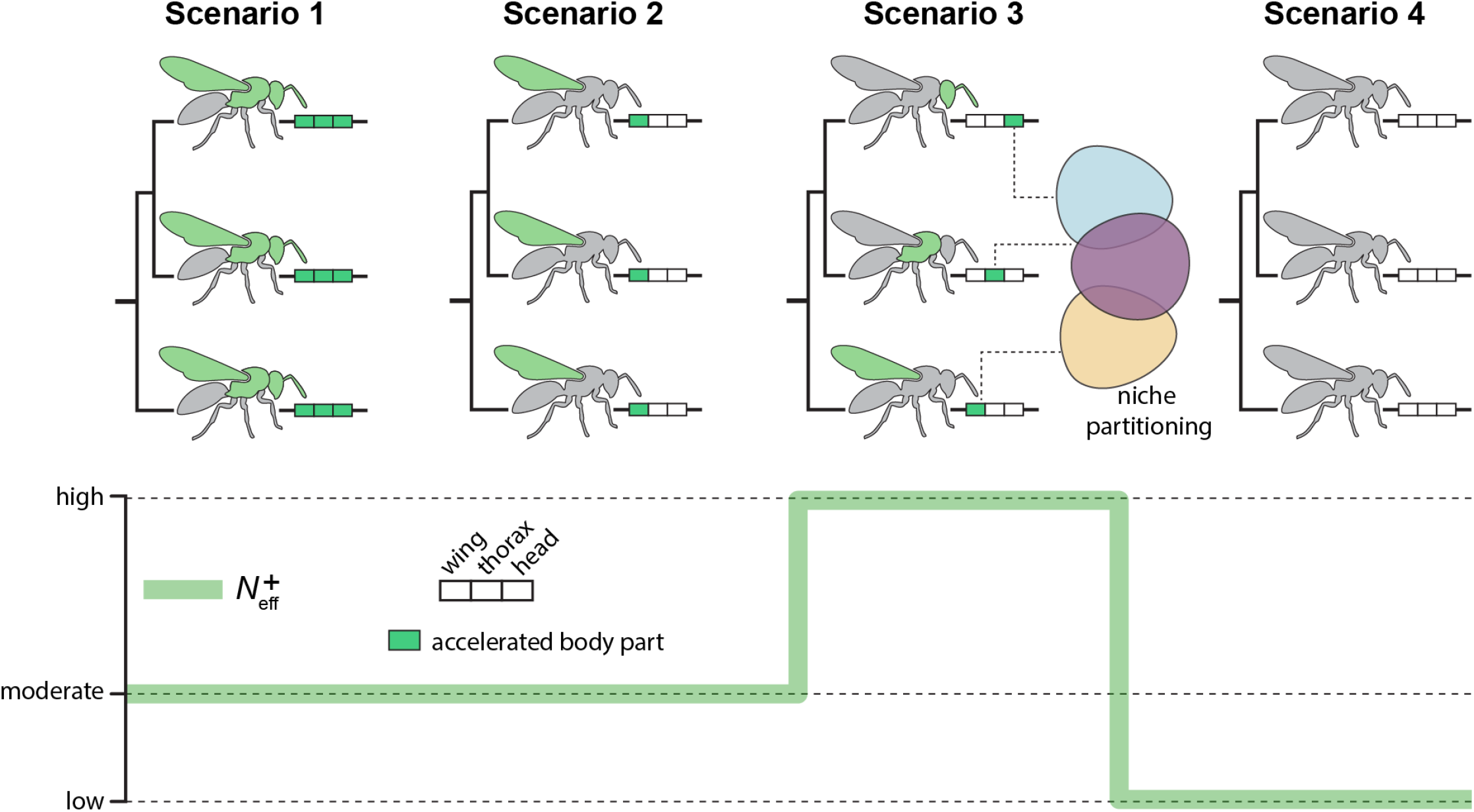
Conceptual illustration of the 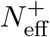 metric. Four hypothetical scenarios illustrating how 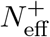 quantifies the diversity of accelerated anatomical systems across lineages, assuming three anatomical regions (head, thorax, and wings). 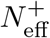 is highest in Scenario 3, where different lineages exhibit acceleration in different anatomical regions. We interpret these alternative phenotypic trajectories as facilitating niche partitioning by expanding the adaptive solutions available to different lineages. 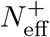 is intermediate when all regions accelerate simultaneously (Scenario 1) or when the same region is accelerated across all lineages (Scenario 2), and lowest when no regions undergo acceleration (Scenario 4).

**Extended Data Figure 5.**
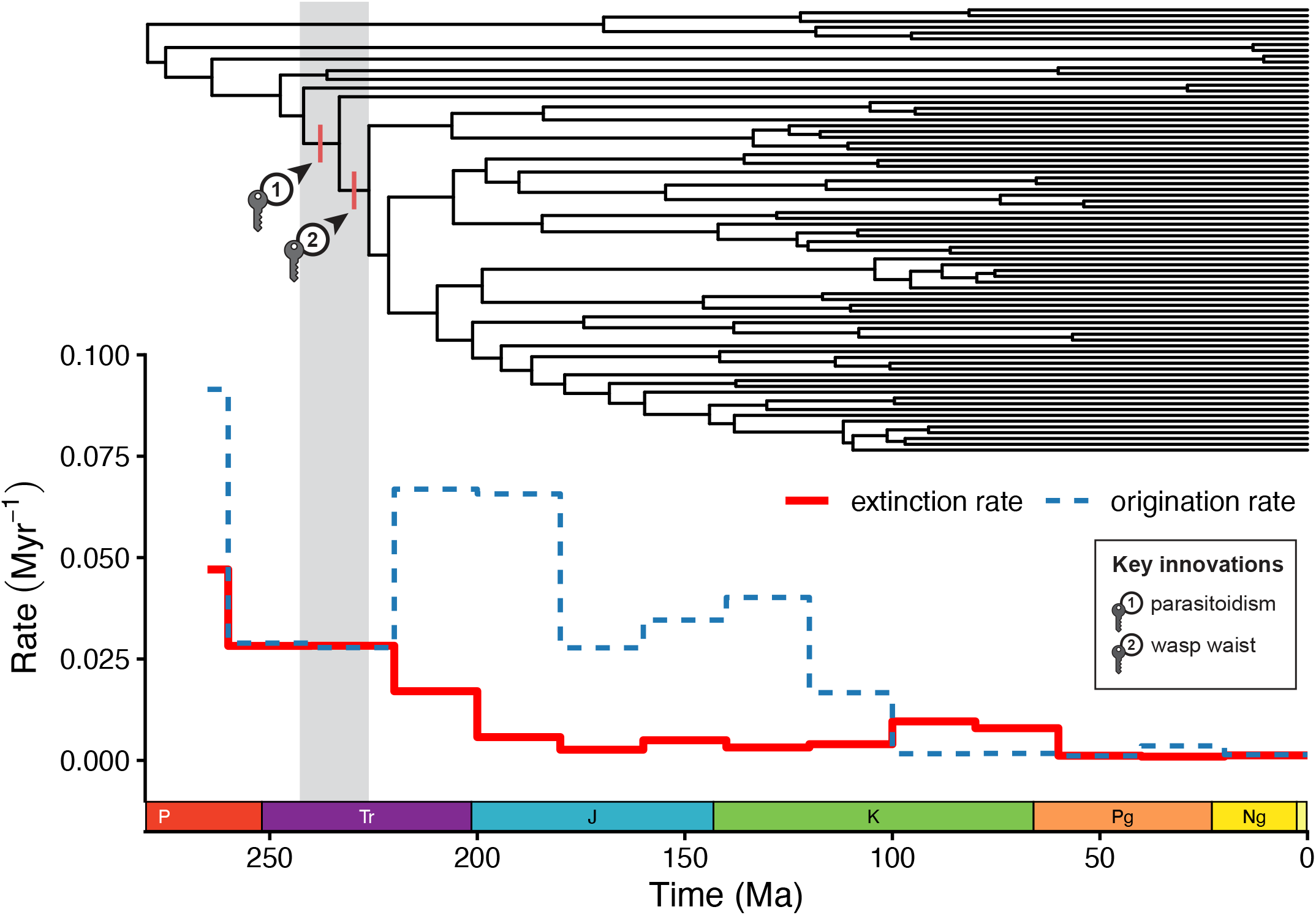
Key innovations and lineage extinction. The origins of larval parasitoidism and the adult wasp waist coincide with elevated lineage extinction rates inferred by PyRate^4^. This pattern is consistent with both innovations representing shifts between major adaptive peaks, with elevated extinction reflecting the traversal of adaptive valleys associated with reduced fitness.

**Extended Data Figure 6.**
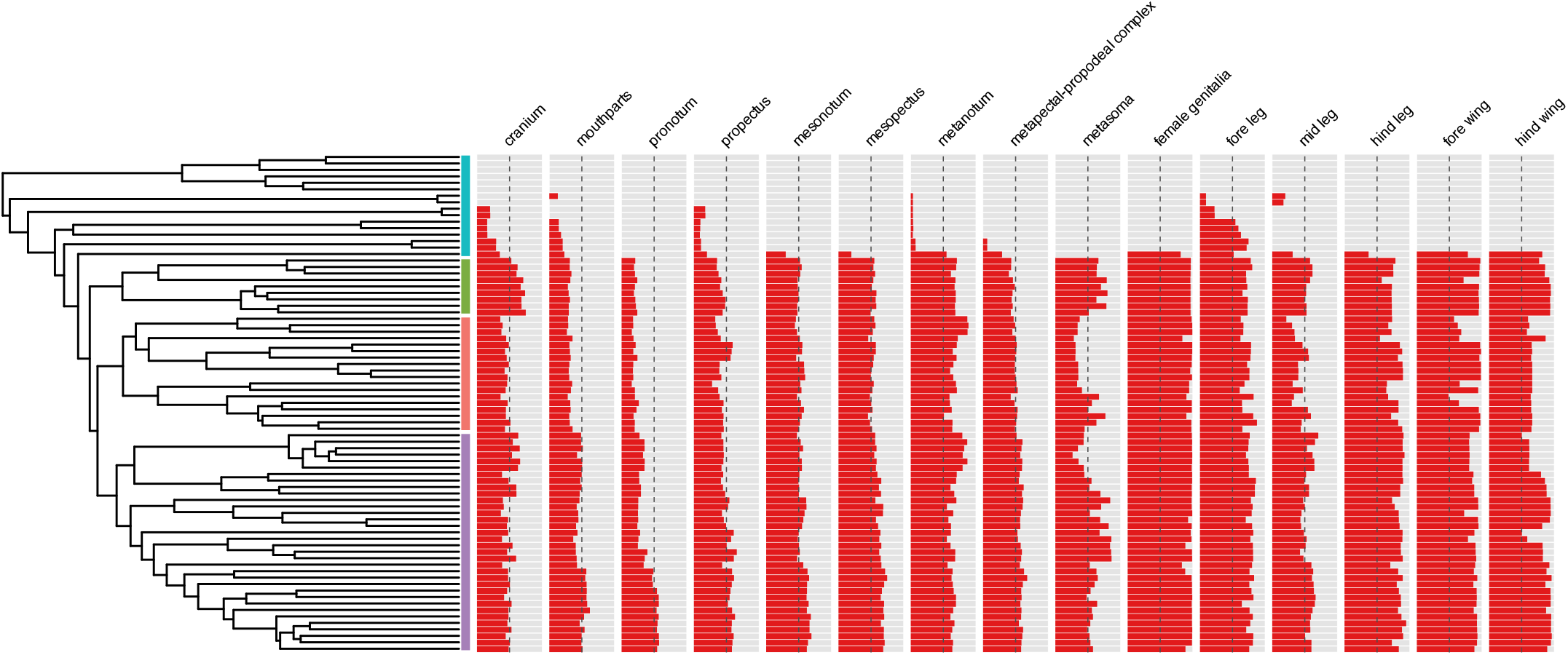
Per-tip proportion of phenotypic changes acquired during accelerated episodes of hymenopteran evolution for each anatomical region. On average, approximately 50% of all phenotypic changes accumulated during accelerated episodes are associated with the rise and diversification of Apocrita. The dashed line indicates the 50% threshold.

